# ADEVO: Proof-of-concept of Adenovirus Directed EVOlution by random peptide display on the fiber knob

**DOI:** 10.1101/2023.11.16.567388

**Authors:** Erwan Sallard, Julian Fischer, Nissai Beaude, Arsalene Affes, Eric Ehrke-Schulz, Wenli Zhang, Adrian Westhaus, Marti Cabanes-Creus, Leszek Lisowski, Zsolt Ruszics, Anja Ehrhardt

## Abstract

Directed evolution of viral vectors involves the generation of randomized libraries followed by artificial selection of improved variants. Directed evolution only yielded limited results in adenovirus vector (AdV) development until now, mainly due to insufficient complexities of randomized libraries.

Clinical applications of AdVs as gene therapy or oncolytic vectors are still hampered by the predetermined tropism of natural types. To overcome this challenge, we hypothesized that the technology of randomized peptide insertions on the capsid surface can be incorporated into the AdV bioengineering toolbox for vector retargeting. Here we developed Adenovirus Directed EVOlution (ADEVO) protocols based on fiber knob peptide display.

As a proof-of-concept, HAdV-C5-derived libraries were constructed following three distinct protocols and selected on A549-DCAR cells that lack the HAdV-C5 primary receptor, with the goal of identifying variants able to infect and lyse these tumor cells more efficiently. All protocols enabled the construction of high complexity libraries with up to 9.6x10^5 unique variants, an approximate 100-fold improvement compared to previously published AdV libraries. After selection, the most enriched variants did not display enhanced infectivity but rather more efficient replication and cell lysis. This warrants investigations into potential unsuspected involvement of the fiber protein in adenovirus replication.

**GRAPHICAL ABSTRACT:** 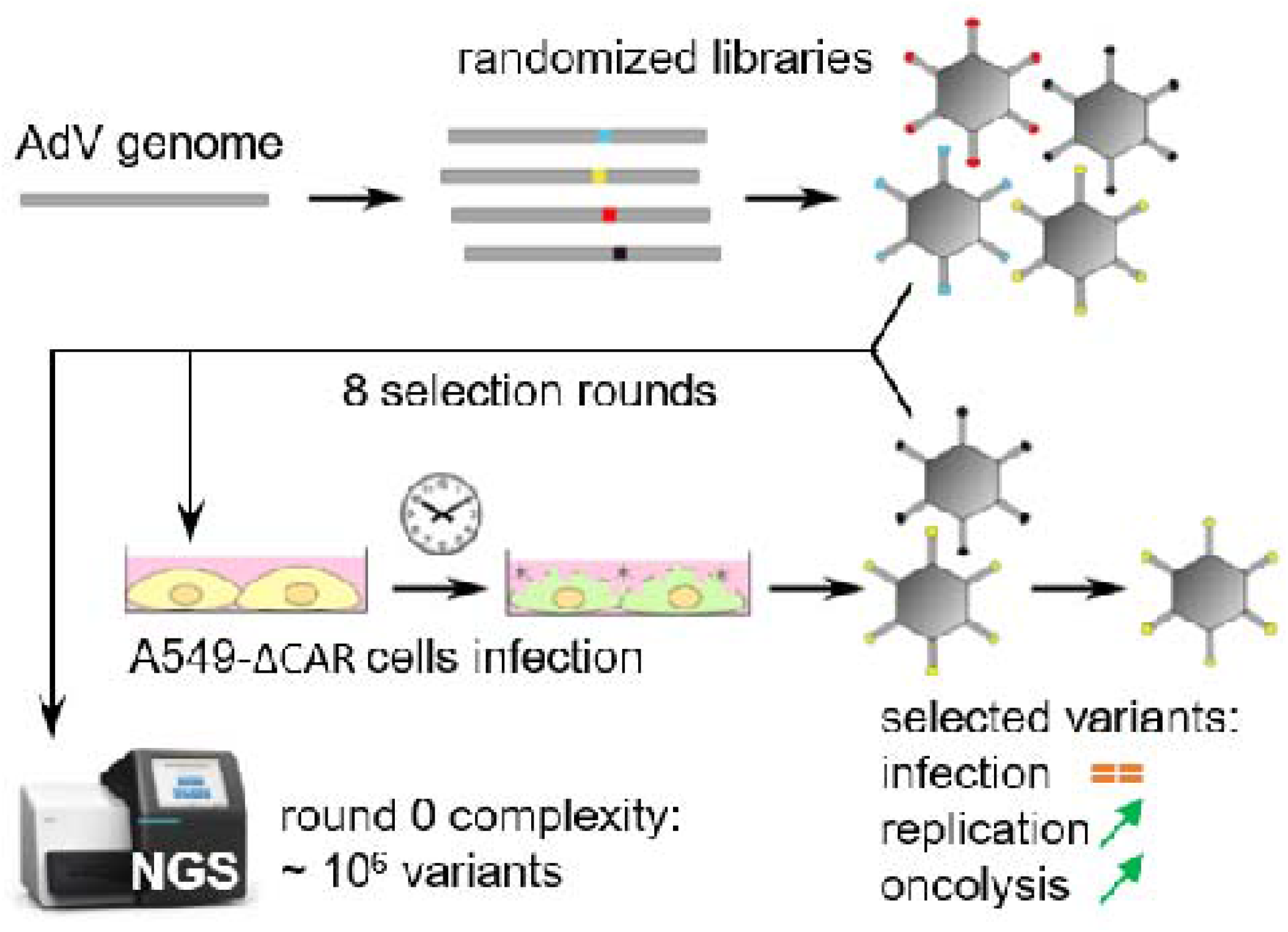

## INTRODUCTION

Adenoviruses are double-stranded DNA viruses with linear genomes of 26 to 45kb and usually cause asymptomatic or benign respiratory or digestive infections in humans (1). There are currently 113 known human adenovirus types (2), grouped in 7 species (A to G). A few of them, most notably human adenovirus type 5 (HAdV-C5) from species C, have been genetically engineered to construct numerous viral vectors for vaccines, gene therapy or oncolytic virotherapy. This makes them the most frequently used type of viral vectors for clinical applications overall (3,4), although in recent years adeno-associated viral vectors (AAVs) have become more popular (5).

Among adenovirus vectors (AdVs), oncolytic adenovirus vectors (OAdVs) are designed to selectively replicate in and lyse tumor cells without harming healthy tissues. Due to their ability to lyse cancer cells, deliver transgenes and locally recruit the immune system, OAdVs can complement already approved therapies such as chemo- and immunotherapies. Dozens of OAdVs are currently undergoing clinical trials (6) and the OAdV Oncorine (H101) was approved by the Chinese Drug Agency (SFDA) for the treatment of nasopharyngeal carcinoma as early as 2005 (7). However, no other OAdV has been approved for clinical use to date.

The knob domain of the fiber protein is responsible for the binding of natural adenovirus types to their cellular primary receptors and thereby contributes to tropism determination. There are only a limited number of adenovirus primary receptors: species A, E, F, C including HAdV-C5, and certain species D types such as HAdV-D9 predominantly use the coxsackievirus and adenovirus receptor (CAR), while other species use primarily CD46, Desmoglein 2 (DSG2) or sialic acid (8). Adenovirus receptors are expressed in numerous cell types, both healthy and tumorous, at varying levels (9). For example, CAR expression levels are high on erythrocytes and in the juvenile brain but very low in several cancer types including pancreatic(10) and colon adenocarcinomas as well as prostate cancer (10,11). Therefore, most potential HAdV-C5-based OAdVs would transduce these tumor types inefficiently while displaying high levels of off-target transduction of healthy tissues, prompting safety issues and further decreasing the number of vector particles (VPs) able to transduce the target tissues. This is particularly concerning given that the cited tumor types still have a very high lethality and medicine would greatly benefit from efficient OAdVs directed against them. Overall, the lack of tropism specificity and the low transduction efficiency are two of the main hindrances in AdV clinical applications, even in case of local delivery (9). Therefore, retargeted AdVs are in high demand.

Several approaches of AdV capsid genetic or chemical engineering have been harnessed to increase the efficiency or specificity of transduction in target tissues. In instances where a target receptor as well as its ligands were known, rational design studies showed that the insertion of short peptides, for example octopeptides, in exposed loops of the fiber knob domain such as the HI loop can lead to successful AdV retargeting (12–14). However, rational design suffers from a low throughput and is not applicable to numerous target cell types for which the optimal vector modifications are not known in advance. This warrants the establishment of new AdV engineering workflows based on complementary principles.

Directed evolution, a concept inspired by the natural processes of random mutation followed by positive selection of the fittest, holds the potential to overcome these shortcomings and substantially accelerate vector bioengineering due to its high throughput and unbiased variant testing. Viral vector directed evolution involves the production of randomized libraries containing thousands of vector variants followed by the selection of the most efficient variants for a given purpose. Random libraries can be produced by 1) insertion of random peptides at specific sites in viral proteins; 2) shuffling of different pre-existing vector types to produce chimeric vectors; and/or 3) random mutagenesis of pre-existing vectors (Figure 1a) (15). The second approach was successfully used with OAdVs in 2008 to obtain the ColoAd1 vector, also known as EnOncoAd (16). ColoAd1 is an Ad3-Ad11 chimera produced by recombination of species B AdV genomes and selected by serial passage of the replication-competent variant library on colon adenocarcinoma cell lines. This vector (under the name Enadenotucirev) and its derivatives have been used in several clinical trials for oncolytic virotherapy (17). However, it remains to this day the only example of successful directed evolution bioengineering in the field of AdV vectorology. Furthermore, random peptide display was used extensively with certain vector systems such as AAVs, yielding for example the AAV2/7m8 vector (18) that is now investigated in several clinical trials of retina gene therapy (NCT03748784, NCT03748784), but despite this success random peptide display was scarcely tested in AdVs. The rarity of AdVs directed evolution studies can be attributed to the technical difficulties of cloning libraries of these large vectors, limiting the variability and size of variant libraries and thus the potency of directed evolution. Indeed, the library complexities (i.e. the number of unique variants present in a library) reported in previous studies of AdV random peptide display reached at best an order of magnitude of only ten thousands (19), while complexities of five millions or more have already been reported with AAVs (20). To put the size of libraries in perspective, saturating a small 10 amino acid domain of a larger protein would require a library in the range of 10^13 (20aa to the power of 10 positions) variants, indicating that current libraries can facilitate examination of only a very small random subset of variants in the space of possibilities. Although these small AdV libraries facilitated the selection of improved OAdVs notably in the AsPc1 pancreatic adenocarcinoma cell line in vitro (21) as well as in vivo in peritoneal tumor xenografts (22), to our knowledge the selected vectors did not make it to clinical trials. Nevertheless, novel high-efficiency genome assembly tools based on Gibson Assembly derivatives may facilitate higher-throughput AdV library cloning and were already successfully used in the context of adenoviruses for rational design applications (23).

**Figure 1.**
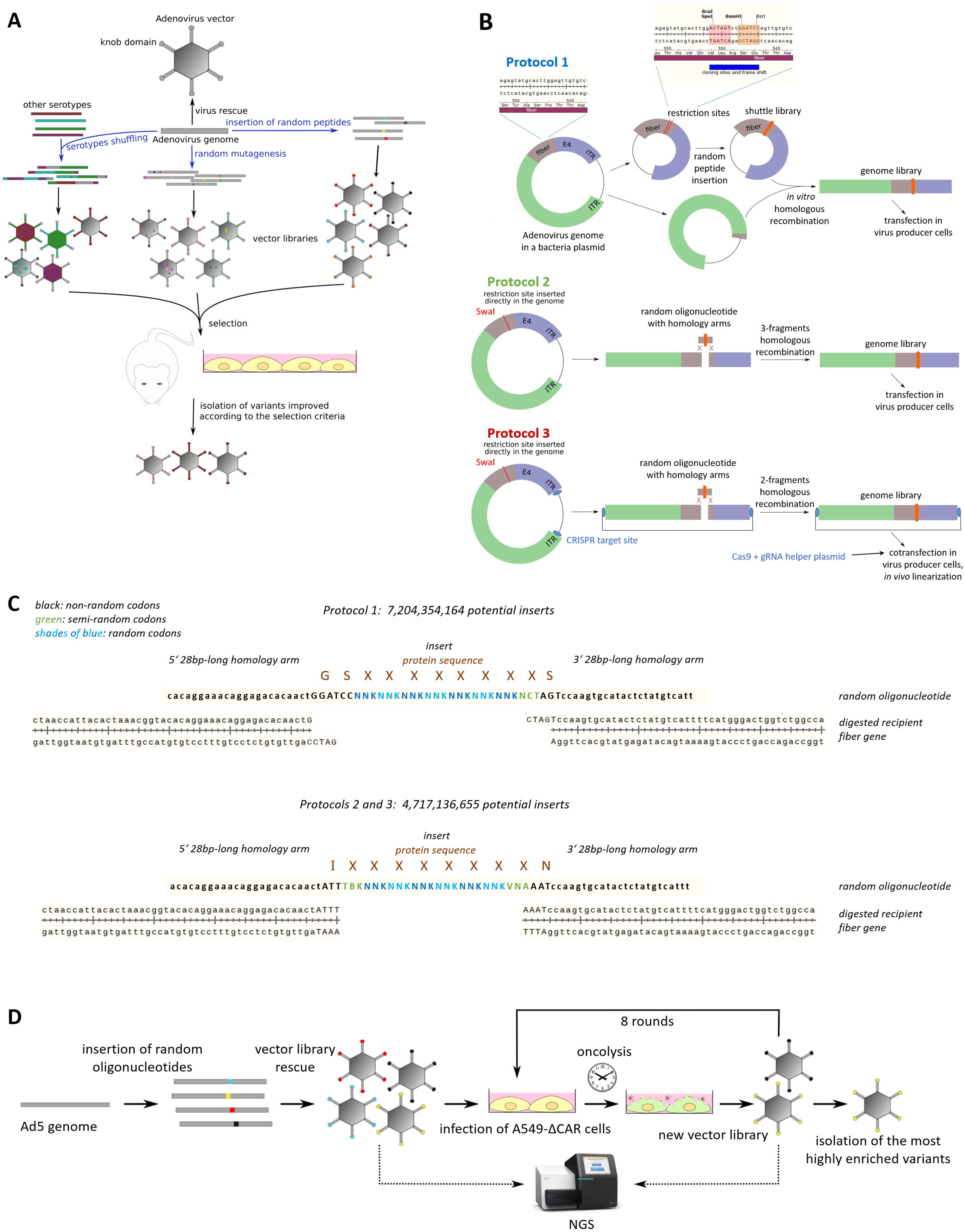
Directed evolution workflows. A: Directed evolution principles. Random libraries can be generated by shuffling of pre-existing vectors, random mutagenesis, or random peptide display. Libraries are subsequently selected in chosen models under stringent conditions to enrich variants optimizing the desired criteria. B: ADEVO protocols for library generation studied in this article. All protocols are based on the insertion of oligonucleotides containing 8 random or semi-random codons in the HI loop of the fiber knob domain. In Protocol 1, libraries are first built in a fiber-carrying shuttle plasmid. The shuttle plasmid is then recombined with a truncated AdV genome to reassemble full insert-carrying AdV genomes. In Protocol 2, libraries are built in a single step by three-fragment homologous recombination between the left and right parts of the AdV genome and the random oligonucleotide. Protocol 3 involves a single-step, two-fragment homologous recombination, and an in vivo Cas9-mediated AdV genome linearization after transfection. C: Sequence details of peptide insertions. Random oligonucleotides used for homologous recombination inside the fiber gene HI loop contain 28 nt-long homology arms matching the fiber (lowercase) and restriction sites (uppercase) sequences of the different protocols, and a central insert of 8 random or semi-random codons. D: Proof-of-concept experiment design. Several libraries were constructed and submitted to 8 rounds of selection on A549-DCAR cells. Full results are available for one protocol 1 library, two protocol 2 libraries and one protocol 3 library. Library quality was assessed by NGS at different steps of the directed evolution workflow. After selection, the most highly enriched variants were isolated and characterized.

Here, we designed and optimized new protocols for adenovirus library generation and assessed them in a proof-of-concept experiment with the goal of selecting improved HAdV-C5-derived oncolytic vectors targeting a CAR-deleted A549 cell line model. Replication-competent AdV libraries were built and selected by serial passaging on target cells, and library complexity and composition was followed by deep sequencing during selection. We show that all three workflows tested facilitated the generation of highly complex libraries comprising hundreds of thousands of unique variants, and the selection of vectors with increased lytic effect in the model cell line.

## MATERIAL AND METHODS

### Cell culture

The A549-□CAR cell line was previously generated by CRISPR-Cas9 deletion of the CXADR gene expressing the CAR receptor (24). A549-WT, A549-DCAR, HeLa, HEK293, Hs578T and MiaPaCa2 cells were cultivated in Dulbecco’s Modified Eagle Medium (DMEM, Pan-Biotech, P04-035911) supplemented with 10% Fetal Bovine Serum (FBS, Pan-Biotech, P40-37500) and Penicillin-Streptomycin (P/S, Pan-Biotech, P06-07100). HCC827 cells were cultivated with Roswell Park Memorial Institute 1640 Medium (RPMI, Pan-Biotech, P04-16500) supplemented with 20% FBS and P/S. All cells were kept at 37 °C under a humidified atmosphere with 5% CO2. Three hours post AdV infection (hpi), cell media was changed and FBS concentration was decreased to 2% (or 5% for HCC827 cells). All cell lines tested negative for mycoplasma infection using the VenorGeM Classic kit (Minerva Biolabs, #11-1025).

### Adenovirus vectors acquisition

PCRs were performed using the Phusion DNA Polymerase (New England Biolabs, #M0530S) following manufacturer’s instructions. All oligonucleotides used in this study are listed in Supplementary Table 1.

Vector genome recombineering was performed as described in Zhang et al. (25). Briefly, selection markers were PCR-amplified using primers with overhangs to obtain homology arms able to recombine with the sequences flanking on both sides the locus to be modified. Selection markers contained an Ampicillin-resistance gene for positive selection and the ccdB toxin for negative selection and were inserted into plasmids containing the vector genome to be modified by homologous recombination in bacteria expressing the λRed recombinase and the mutated ccdA antitoxin (GB05-red with GyrA mutation of Arg462Cys). In a second step, the selection marker was removed and replaced by the desired sequence flanked by homology arms, coming either from an oligonucleotide in the case of point mutations or from PCR amplification when larger sequences needed to be inserted. This homologous recombination was performed in bacteria expressing the RecET recombinase system (26).

Protocol 1 assembly backbone was obtained by replacing all HAdV-C5 sequences between the fiber gene and the right ITR with a PacI restriction site. The deleted sequences plus a 20nt-long homology arm in 5’ of the fiber gene were subcloned into a pJet plasmid (CloneJET PCR cloning kit, ThermoFischer, K1231) and flanked by BstBI restriction sites to obtain the shuttle plasmid. BamHI-HF (New England Biolabs, #R3136S) and SpeI-HF sites (New England Biolabs, #R3133S), separated by two nucleotides as a frameshift to avoid rescue of undesired parental vectors during library production, were inserted between the codons 546 and 547 of the fiber gene, corresponding to the HI loop position (19) (Figure 1b). To this end, the whole shuttle plasmid was PCR-amplified with overlapping primers containing the desired insert and recircularised by NEBuilder (New England Biolabs, #E2621L) recombination.

Protocol 2 assembly backbone was obtained by insertion of a SwaI restriction site between the codons 546 and 547 of the fiber gene. This introduced a frameshift and stop codon aimed to avoid rescue of undesired parental vectors during library production. Protocol 3 assembly backbone was derived from the protocol 2 assembly backbone by replacing the bacterial plasmid backbone so that the final product does not contain SwaI restriction sites except the one in the fiber gene, and contains the int5 gRNA sequence (tatattattgatgatgCCTC) followed by a CGG PAM at the end of both ITRs, as described in Riedl et al. (27), to facilitate CRISPR-mediated vector genome linearization (Figure 1b).

To purify the ADEVO-selected variants, oligonucleotides containing their inserts and surrounding sequences were reassembled into the Protocol 2 assembly backbone using the same methods as for random oligonucleotides during Protocol 2 library production. The re-cloned ADEVO-selected variants, as well as the HAdV-C5-DFiber vector and HAdV-C5-WT control vector, were rescued and purified as described in Jäger et al. (28) and titrated by optical density measurements as well as quantitative PCR (qPCR) using a CFX96 Real-Time System machine (BioRad) and the my-Budget 5x EvaGreen qPCR-Mix II (Bio-Budget, #80-5901000) following the manufacturer’s instructions.

The luciferase-expressing HAdV-C5 vector has already been described in Zhang et al. (25).

### Cross-packaging assay

Adherent HEK293 cells grown to 80% confluency were transfected using the jetOPTIMUS reagents (Polyplus) following manufacturer’s instructions with model mini-libraries consisting in SwaI-linearized HAdV-C5-WT and HAdV-C5-DFiber genomes at a 1:1 molar ratio, with varying total quantities of transfected genomes per cell. Cells and media were harvested when cytopathic effect (CPE) was observed. The lysates underwent four cycles of freeze and thaw followed by benzonase (Emprove expert benzonase endonuclease, Merck #101695) digestion of non-encapsidated DNA at 37 °C for 30 min with gentle shaking using 125 U of enzyme per mL of lysate. Progeny vector particles (VP) were titrated by qPCR from lysate dilutions, while subconfluent HEK293 cells were infected using 40% of total lysate volume per well of 6 well plate. At 3 hpi, infected cells were washed five times to eliminate non-internalized VPs and DNA was extracted using the NucleoSpin Tissue kit (Macherey-Nagel #740952-250). Internalized vector genomes were titrated by qPCR using a forward primer in the U exon, a reverse primer within the fiber gene to titrate HAdV-C5-WT genomes, and a reverse primer downstream of the fiber gene (Supplementary Table 1) to titrate HAdV-C5-DFiber genomes. The cross-packaging rate was calculated as the proportion of HAdV-C5-DFiber genomes among all internalized vector genomes.

### Random library production

Shuttle plasmids and Protocols 1, 2 and 3 assembly backbones were digested for 24 hours using respectively the BamHI-HF and SpeI-HF; PacI; PacI then SwaI (24 hours each); and SwaI restriction enzymes (New England Biolabs), then purified with phenol – chloroform - isoamyl alcohol (Roth, #A156.3), precipitated with ethanol and resuspended in pure water.

The first phase of Protocol 1 consists in constructing a shuttle library following methods already established for random peptide display on AAVs (29). NEBuilder homologous recombination was performed using 225 fmol of BamHI-HF-SpeI-HF-digested shuttle plasmid and 2.25 pmol of single stranded oligonucleotides containing 28 nt-long homology arms to each end of the BamHI-HF-SpeI-HF-digested shuttle plasmid and a central NNK7 motif coding for a random heptapeptide with lower redundancy (Figure 1c). The recombination products were purified with phenol – chloroform - isoamyl alcohol, precipitated with ethanol and resuspended in pure water, then electroporated in competent E. coli. Following 1 h recovery in pure LB media, the electroporated bacteria were cultivated overnight in 100 mL of LB media supplemented with 50 µg/mL ampicillin. The shuttle library was extracted using the ZymoPure II plasmid midiprep kit (Zymo Research, #D4201) and digested with BstBI (New England Biolabs) for 24 hours, then purified with phenol – chloroform - isoamyl alcohol, precipitated with ethanol and resuspended in pure water.

Protocol 1 genome libraries were reassembled by NEBuilder homologous recombination using 375 fmol of the BstBI-linearized shuttle library and 375 fmol of the PacI-linearized Protocol 1 assembly backbone. Protocols 2 (respectively Protocol 3) genome library reassembly was performed in one step using 4.5 pmol (respectively 2.25 pmol) random oligonucleotides and 225 fmol of PacI-SwaI-digested (respectively SwaI-digested) assembly backbone for three-fragments (respectively two-fragments) NEBuilder homologous recombination. The recombination products were purified with phenol – chloroform - isoamyl alcohol, precipitated with ethanol and resuspended in pure water.

For all protocols, round 0 libraries were obtained by transfecting 4.8 µg of genome library in 30 million 80% confluent low passage number HEK293 cells (corresponding to approximately 4,000 reassembled genomes per cell if the reassembly was 100% efficient) using the jetOPTIMUS reagents (Polyplus, #101000006) following manufacturer’s instructions. In the case of Protocol 3, in vivo vector genome linearization was facilitated by cotransfection of 9.6µg of pAR-Int5-Cas9-Amp helper plasmid as described in Riedl et al. (27). For all libraries, CPE was observed 3 days post transfection and cells and media were harvested. The lysates were frozen and thawed four times then used for selection.

### Vector selection

At each selection round, in order to determine library transduction titers, A549-DCAR cells cultivated in 24 well plates were transduced using 50, 10 or 2 µL of the lysate obtained from the previous selection round. At 3 hpi, cells were washed five times to eliminate non-internalized VPs and harvested. DNA was extracted using the Macherey-Nagel Nucleospin Tissue mini kit and internalized vector genomes were titrated by qPCR. Subsequently, ten million confluent A549-DCAR cells were infected by the desired libraries at 0.5 transducing units per cells. Media was changed at 3 hpi and cells and media were harvested 4 days post infection (dpi) and frozen and thawed four times. This lysate would then be used for the next selection round. Eight rounds of selection were performed.

### Next generation sequencing (NGS)

Library purity and complexity was assessed by NGS after round 0, round 8 and optionally round 4. Aliquots consisting of 0.25% of the total lysate volume were taken from the desired libraries before freeze and thaw cycles had been applied and were processed for NGS. Cells were pelleted by centrifugation at 6,000 g for 1 min. The pellet was resuspended in benzonase buffer (50 mM Tris-HCl pH=8.5, 2 mM MgCl2) and frozen and thawed four times. Both the supernatant and the resuspended pellet were incubated with 1 U/µL of benzonase (Emprove expert benzonase endonuclease, Merck #101695) at 37 °C for 1 h with gentle shaking to digest non-encapsidated DNA. In our hands, this treatment facilitated the digestion of >99.9% of non-encapsidated DNA while leaving encapsidated viral DNA untouched (Supplementary Figure 4). Benzonase was then inactivated and vector genomes were released by incubation in TE buffer with 0.5 mg/mL proteinase K, 5mM EDTA and 0.5% SDS for 3 h at 56 °C with gentle shaking. Proteinase K was heat-inactivated at 80 °C for 10 min, then the cell and media samples were regrouped and vector DNA was purified with phenol – chloroform - isoamyl alcohol, precipitated with ethanol and resuspended in pure water. Variants inserts and surrounding regions were PCR amplified using the entirety of purified DNA, PrimeStar max DNA polymerase (Takara Bio, #R045A) and barcoded primers (Supplementary Table 1) with a total of 30 cycles. The 174 nt-long (Protocol 1) or 171 nt-long (Protocols 2 and 3) amplicons were separated from primers and free nucleotides by electrophoresis in a 2% agarose gel then gel extracted using the MyBudget double pure kit (Bio-Budget, #55-3000) and sequenced by Eurofins Genomics under the NGSelect amplicon adaptor ligation package. Up to 6 library aliquots were pooled together for sequencing and separated during bioinformatic analysis thanks to their different barcodes. Primer barcodes were 4 nt-long and had at least three bases different from each other in order to minimise the risk of misidentification.

Three aliquots of the round 0 libraries were processed and sequenced to enable statistical estimation of library complexity.

NGS output reads were subjected to quality control in order to exclude library contaminants including insertless parental genomes and, as well as PCR and sequencing errors. Criteria for read exclusion were lack of read pairing within the same library aliquot, incorrect read length, lack of appropriate primer barcode, mutations in the sequences flanking inserts on either side, NGS quality score inferior to 30 on the insert sequence, and mismatch between paired reads. Reads with unexpected sequences at the two nucleotide positions directly preceding the insert or the two nucleotides positions directly following it were considered as having incorrect flanking sequences, since it meant that the insert could not be identified.

Wild-type fiber sequences and parental fiber sequences (corresponding to the assembly backbones) were counted as library contaminants. Inserts that passed all quality controls were translated to obtain their amino acid sequences. Unique insert sequences were numbered and listed alongside their read counts in the corresponding library aliquots.

NGS variant calling data was analyzed on python using the packages pandas (30), numpy (31) and Bio.Seq (32). The package matplotlib.pyplot (33) was used for graphical representations.

### Infectivity assays

Cells cultivated in 24 well plates were infected by 10 vpc of ADEVO variants or HAdV-C5-WT. At 3 hpi, cells were washed five times to eliminate non-internalized VPs and harvested. DNA was extracted using the Macherey-Nagel Nucleospin Tissue mini kit and internalized vector genomes were titrated by qPCR.

### Luciferase assays

Cells cultivated in 96-well plates were infected by 20 vpc of luciferase-expressing HAdV-C5 vectors. Media was changed at 3 hpi and luciferase luminescence was measured at 24 hpi using the Nano-Glo® Luciferase Assay (Promega, Madison, USA, #N1130) kit, a TECAN infinite f plex plate reader and black 96-well luciferase plates (Thermo Fisher Scientific Nunc A/S).

### Replication assays

Cells cultivated in 24-well plates were infected by 10 vpc of the chosen vectors. Media was changed at 3 hpi. At the designated time points, cells were either washed five times and harvested for titration of intracellular vector genomes performed as for infectivity assays, or media was collected and replaced to titrate excreted progeny vector particles. To this aim, 250 µL of media were treated with 125 U/mL benzonase at 37 °C for 30 min with gentle shaking to digest non-encapsidated DNA. Benzonase was then inactivated and vector genomes were released by incubation in TE buffer with 0.5 mg/mL proteinase K, 5 mM EDTA and 0.5% SDS for 3 h at 56 °C with gentle shaking. Proteinase K was heat-inactivated at 80 °C for 10 min, then vector DNA was purified with phenol – chloroform - isoamyl alcohol, precipitated with ethanol, resuspended in pure water and titrated by qPCR.

### Oncolysis assays

Cells cultivated in 96-well plates were infected by 1, 3, 10, 30, 100, or 300vpc of ADEVO variants or HAdV-C5-WT and media was changed at 3 hpi. When CPE was observed for at least one vector in the 1 vpc condition, viability measurements were performed for the 1 vpc and 3 vpc conditions of all vectors and two wells of uninfected cells with the Cell Counting Kit 8 (CCK8, Sigma-Aldrich #96992). Subsequently, all cells were washed twice with PBS, fixed for 15 min at room temperature (RT) in PBS + 2% paraformaldehyde, and washed twice with PBS. Live cells were stained for 10 min at RT in methanol + 2% crystal violet. Pictures were taken after thorough washing and overnight drying.

### Statistics

Total library complexity was estimated using Anne Chao’s simple capture-recapture Mh-type model (34) after counting the overlap of variant composition between at three library aliquots. If more than three aliquots had been sequenced for a library, the average of the complexity estimates computed for each possible set of three aliquots was considered. More details on statistical modeling are reported in Supplementary Figure 2.

## RESULTS

### ADEVO protocols facilitate the production of highly complex random libraries

We designed three protocols for Adenovirus vector Directed EVOlution (ADEVO). Protocol 1 was inspired by AAV random library generation methods (29): HAdV-C5 fiber, E4 region and right ITR were subcloned into a 7 kb-long shuttle plasmid, corresponding to the size of typical AAV plasmids. BamHI and SpeI restriction sites were inserted into the fiber HI loop region (Figure 1b) so that random oligonucleotides containing seven random codons and one semi-random codon could be inserted into the shuttle plasmid by high throughput homologous recombination (Figure 1c). Obtained shuttle library plasmids could then be linearized and recombined with plasmids carrying all HAdV-C5 sequences absent from the shuttle plasmid, namely those from the left ITR to the U exon, in order to reassemble functional HAdV-C5-derived genomes carrying random inserts in their fiber gene. Reassembled genome libraries are to be transfected in producer cells to rescue the round 0 vector libraries. To minimize the risk of library contamination by insert-less religated vectors, two nucleotides were inserted in the shuttle plasmid between the BamHI and SpeI restriction sites so that insert-less shuttle plasmid would give rise after genome reassembly to non-functional genomes with a frameshift in the fiber gene.

Alternatively, ADEVO Protocol 2 consists of a single-step library generation whereby a SwaI restriction site, absent from the HAdV-C5-WT genome, was inserted at the chosen position in the fiber HI loop region (Figure 1b). After digestion of the assembly backbone, three-fragment homologous recombination was conducted between a random oligonucleotide and the two backbone genome fragments left and right of the SwaI site (Figure 1c). Here, random oligonucleotides contained six random and two semi-random codons to minimize differences in the maximal theoretical library complexity between both protocols since the flanking sequences were different from Protocol 1. Like in Protocol 1, reassembled Protocol 2 genome libraries can be transfected into producer cells to rescue round 0 vector libraries, then undergo selection. Likewise, the SwaI restriction site introduced a frameshift and a stop codon which ensured that parental assembly backbones would not contaminate the vector libraries.

AdV rescue was reported to be less efficient after transfection of already linearized AdV genomes than after transfection of circular plasmids carrying AdV genomes followed by in-cell genome linearization by CRISPR-Cas9 (27). We thus developed Protocol 3 derived from Protocol 2 in order to use this improved transfection method. To do so, we replaced the SwaI and PacI restriction sites in the bacterial backbone of the parental AdV genome plasmid with CRISPR-Cas9 target sites so that the genome reassembly products remain circular and can be linearized in producer cells after co-transfection with helper plasmids expressing Cas9 and the specific gRNA (Figure 1b).

Each protocol was optimised with the goals of maximizing purity and yield at each step, and of minimizing cross-packaging. Cross-packaging occurs when a capsid carries a genome encoding for a different capsid sequence (35). In directed evolution experiments, this phenotype-genotype discrepancy can entail selection failure, as selected capsid properties are lost at the next vector generation. We quantified for the first time Adenovirus cross-packaging, in the context of vector rescue after AdV genome transfection, corresponding to library production setting. We detected dose-dependent rates of cross-packaging and found that transfection of 5,000 AdV genomes per cell or less facilitated high titer vector rescue while keeping cross-packaging rates below 3% (Supplementary Figure 1). We chose to use these conditions for library generation.

Next, we designed a proof-of-concept experiment with the aim of comparing the ADEVO protocols and establishing the feasibility and usefulness of directed evolution for AdVs. Since improved oncolytic vectors are in high demand for lung cancer (36), we used A549-DCAR cells, derived from the A549 lung adenocarcinoma model cell line by knock-out of the primary receptor of wild-type HAdV-C5 (24), as target cells for our libraries. It could be expected that highly complex libraries of fiber-modified HAdV-C5 variants would contain vectors able to efficiently infect lung cancer cells through CAR-independent mechanisms. To select for such variants retargeted to A549-DCAR cells, cells were infected with AdV libraries and following replication, progeny vectors were harvested as a new library for subsequent selection rounds. Overall, eight rounds of iterative selection were performed per ADEVO library, and the final enriched variants were identified by Next-Generation Sequencing (NGS; Figure 1d). Finally, the efficiency of ADEVO protocols would be assessed depending on round 0 library purity and complexity, and the oncolysis ability of the selected variants in A549-DCAR cells.

To estimate the total library complexity at round 0, we assessed several statistical models of capture-recapture (Supplementary Figure 2a). These models were developed with the aim of estimating animal populations by capturing, marking and releasing animals, then capturing a new “population aliquot” and counting how many of the animals had already been identified in the first sampling. Here we adapted these models by sequencing several library aliquots, considering each insert peptide sequence as an individual “animal” and counting in how many aliquots it was identified. We used so-called Mh-type models, which take into account that variants have different probabilities of being sampled in none, one or several aliquots due to abundance differences; and Mth-type models, which in addition account for varying aliquot sizes. We found Anne Chao’s estimator (34) to be the most consistent for our data and used it henceforth. Of note, it is a conservative estimator designed to indicate the lower bound of the confidence interval of population size.

A total of four libraries underwent the entire directed evolution process: one from Protocol 1, two from Protocol 2 and one from Protocol 3. Their estimated complexity at round 0 reached over 963,000 for one Protocol 2 library and 245,000 for the other, while the Protocol 1 library counted over 874,000 unique variants estimated and the Protocol 3 library over 166,000 (Figure 2a). The observed complexity in the sequenced library aliquots was respectively over 253,000, 88,000, 334,000 and 42,000 (Supplementary Figure 2b). Protocols 2 and 3 round 0 libraries showed already large differences in individual variants relative abundance (Figure 2b), while the Protocol 1 library variants abundance distribution was flatter. The library purity ranged between 94% and 98% for Protocols 2 and 3 libraries while reaching well over 99% for the Protocol 1 library (Figure 2c).

**Figure 2.**
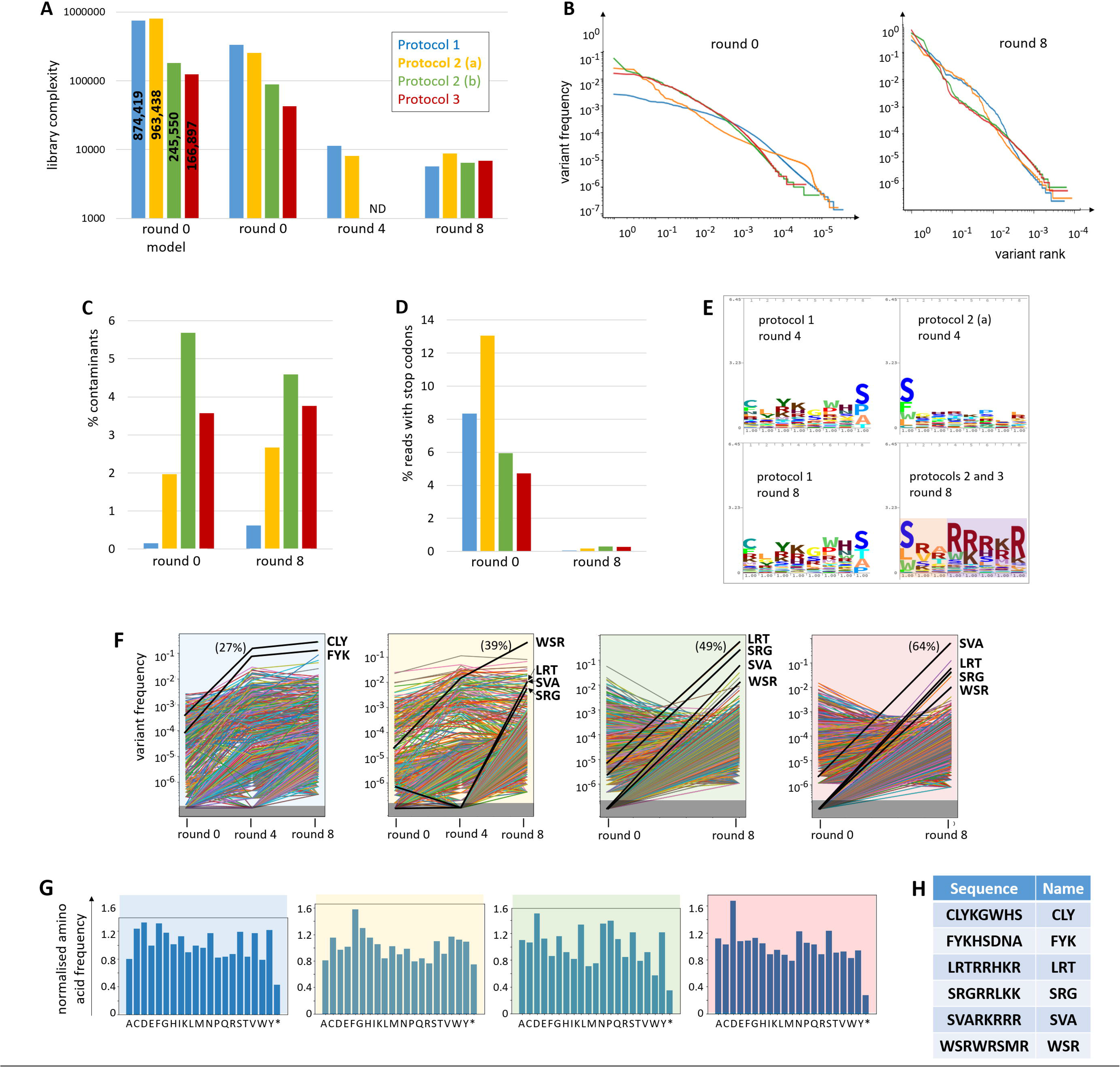
Libraries complexity, purity and composition before and after selection. One library assembled following Protocol 1, two following Protocol 2 (termed „a“ and „b“) and one following Protocol 3 underwent eight selection rounds on A549-DCAR cells. All libraries were sequenced by NGS after rounds 0 and 8, as well as libraries 1 and 2(a) after round4. A: Statistical modeling estimation of total round 0 library complexity („round 0 model“), or number of variants identified by NGS („round 0“, „round 4“, „round 8“). B: Proportion of round 0 (left) and round 8 (right) libraries occupied by each variant, ranked in decreasing abundance. C: Contaminants (parental genomes, inserts with incorrect length, variants with mutations outside the insert, etc) identified by NGS remained rare in the libraries. D: Percentage of all NGS reads containing stop codons in the insert sequence. The loss of inserts with stop codons accross selection rounds indicates that non-functional variants were counter-selected. E: Logo profiles of insert sequences in selected libraries. The aminoacid sequences of the insert random and semi-random positions of the 100 most frequent variants of the considered library were aligned, with a number of repeats per variant proportional to their detected frequency in the library. For round 8, the protocol 2 and 3 libraries were combined given that they have the same insert flanking sequences (SwaI restriction site) and the most highly enriched variants of these libraries largely overlapped. After 8 rounds of selection, a distinct pattern emerges in protocol 2 and 3 libraries: mostly hydrophobic residues in the first three positions, strong predominance of positively charged residues in the last five. F: Evolution of variant frequencies within libraries during selection. The most highly enriched variants (bold lines) were chosen for purification. In brackets: proportion of the library occupied by the dominant variant after selection. Black rectangle: undetected variants. Only variants detected at round 8 are represented. G: Frequencies of each amino acid at selection round 0 within the fully random insert sequence normalised on theoretical expected frequencies show limited initial sequence bias. H: Most highly enriched variants in the different libraries at the end of 8 selection rounds. They will be refered to in the text by the name indicated in the right column.

### Selection of improved oncolytic Adenovirus vectors

All four libraries were submitted to 8 rounds of selection each on A549-DCAR cells. As anticipated, library complexity decreased (Figure 2a). Variants with stop codons in their inserts were virtually eliminated from the libraries (Figure 2d), showing that non-functional variants had been counter-selected. Meanwhile, library purity remained approximately constant across selection rounds (Figure 2c). The dominant variant of each library reached 27% to 64% abundance within the entire library (Figures 2b and f). The abundance distribution had become substantially steeper than at round 0. A limited number of variants had been highly enriched; several hundreds of variants subsisted at low frequency, representing presumably bystander variants that had been able to infect and replicate in the target cells but at a disadvantage compared to the enriched variants; finally, thousands of variants had been eliminated from the libraries during selection (Figure 2b).

A distinctive insert sequence pattern could be observed in Protocols 2 and 3 libraries at selection round 8, with the first three semi-random and random positions predominantly occupied by hydrophobic amino acids, contrasting with the strong enrichment of positively charged amino acids in the last five positions (Figure 2e). This pattern was not clearly detectable at round 4 (Figure 2e) and the amino acid distribution at round 0 showed little enrichment or impoverishment in any amino acid (Figure 2g), indicating that the pattern was not the result of initial biases in library construction. The Protocol 1 library did not display distinctive insert sequence motifs, which may be attributed to the different insert flanking sequences compared to Protocols 2 and 3.

The most highly enriched variants in the different libraries were identified (Figure 2h), re-cloned and purified. Interestingly, the sets of most highly enriched variants from Protocols 2 and 3 libraries largely overlapped (Figure 2f).

The purified variants were compared with each other and with wild-type HAdV-C5 for their infectivity, replication speed and oncolysis strength in target and non-target cell lines. Contrary to expectations, none of the variants displayed strongly increased internalization in A549-DCAR cells compared to HAdV-C5-WT, which was relatively efficient at transducing this target cell line compared to other cell lines (Figure 3a). The internalization rate in A549-DCAR cells was proportional to that in A549-WT cells, and except for the FYK variant correlated to that in unrelated HeLa cells, indicating that the variants were not retargeted to new receptors. We confirmed that HAdV-C5-WT internalization in A549-DCAR cells corresponded not only to abortive particle uptake but also productive transduction by studying transgene expression of a luciferase-expressing vector with HAdV-C5-WT capsid, which confirmed that the infectivity of HAdV-C5-WT in A549-DCAR cells reaches around 20% of the level observed in A549-WT cells (Figure 3b). This indicates that the selective pressure on the cell entry step was lower than expected, and suggests that the variants were selected for other features of their life cycle.

**Figure 3.**
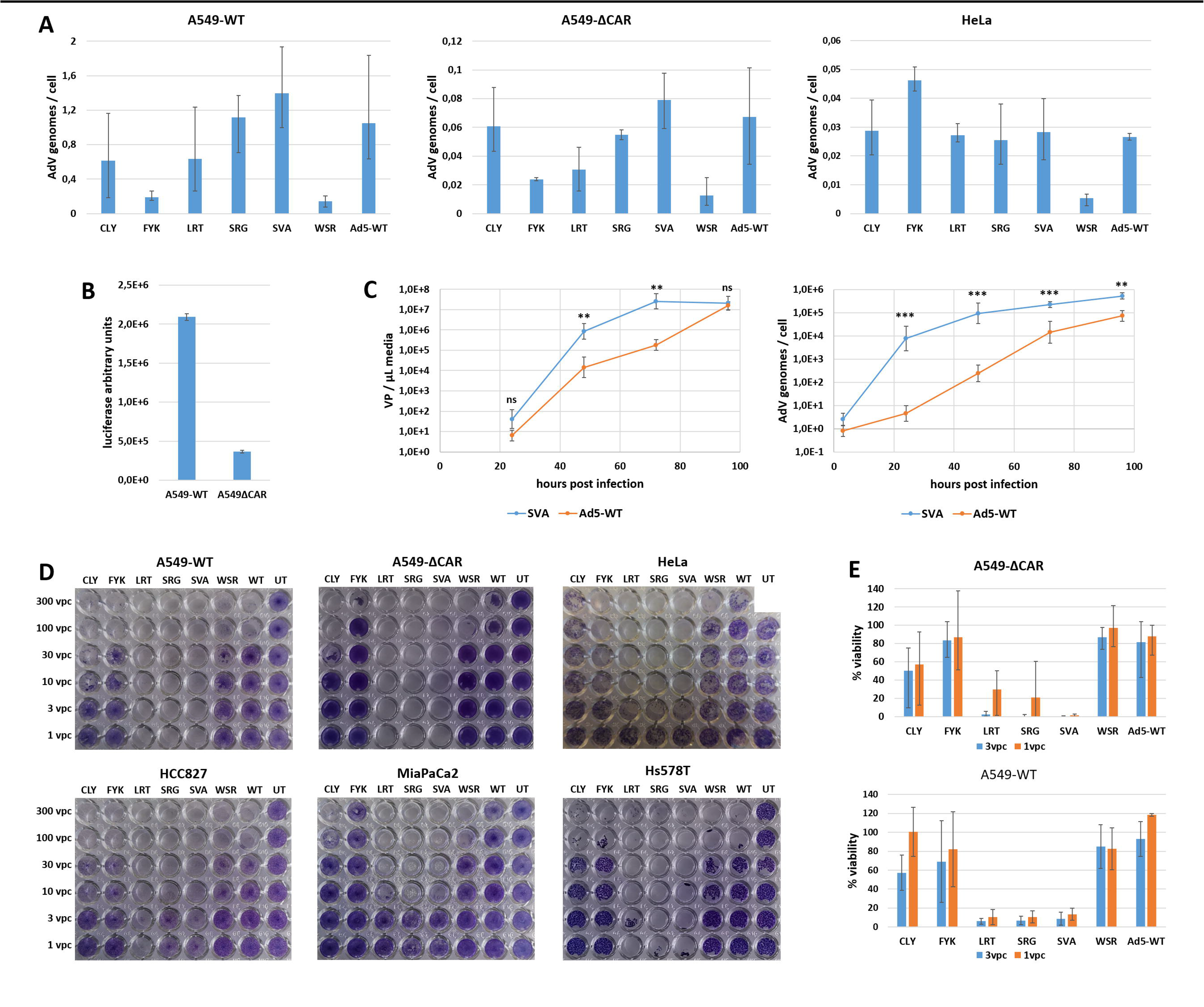
ADEVO-selected variants show improved cell lysis and DNA replication but not increased infectivity. A: ADEVO-selected variants do not display modified tropsim. A549-WT, A549-DCAR and HeLa cells were infected by 10 vector particles per cell (vpc) of HAdV-C5-WT or ADEVO-selected variants. At 3 hours post infection (hpi), internalized vector genomes were quantified. ADEVO-selected variants do not display preferential infection of the A549-DCAR cells in which they were selected compared to other cell lines, while HAdV-C5-WT infected A549-DCAR cells efficiently. N=6, two independent experiment repeats, error bars: maximum-minimum. B: A549-DCAR cells are permissive to non-modified HAdV-C5 infection. A549-DCAR and A549-WT cells were infected at 20 vpc by a luciferase-expressing HAdV-C5 vector. Luciferase luminescence was quantified at 24 hpi, showing relatively strong viral transgene expression in A549-DCAR cells. N=2. C: The SVA variant replicates faster than HAdV-C5-WT. A549-WT cells were infected with 10vpc of HAdV-C5-WT or SVA variant. At regular intervals, culture supernatant was collected and excreted VPs were quantified. Alternatively, cells were harvested, washed thoroughly, and intracellular AdV genomes were quantified. N=6, two independent repeats, data points: geometric mean ± standard deviation. SVA and HAdV-C5-WT titers at each time point were normalised on the respective internalised genome titers at 3 hpi and compared by Welch T test (Mann Whitney U tests confirmed the significance in all cases where T test indicated it). ns: p > 0.05. **: 0.01 > p > 0.001. ***: 0.001 > p. Two-sided p-values. D: The LRT, SRG and SVA variants lyse efficiently a wide array of cancer cell lines. Cancer cell lines were infected by HAdV-C5-WT or ADEVO-selected variants at various multiplicities of infection ranging from 1 to 300 vpc. Several days later (3 dpi for A549-WT cells, 5 dpi for HCC827 and Hs578T cells and 6 dpi for A549-DCAR, HeLa and MiaPaCa2 cells), live cells were stained with crystal violet. UT: untransduced cells; WT: HAdV-C5-WT. Two to four experiment repeats were performed, representative pictures are shown. E: CCK8 quantification of infected A549-DCAR and A549-WT cells viability. Prior to crystal violet staining (Figure 3d), the viability of cells infected with 3 vpc or 1 vpc was quantified and normalized on the viability of uninfected cells cultivated in the same conditions. N=3, three independent experiment repeats, error bars: maximum-minimum.

We indeed observed increased cell lysis of the selected variants, and in particular the LRT, SVA and SRG variants, compared to HAdV-C5-WT (Figures 3d and e). This effect occurred not only in A549-DCAR cells, but also in all other tested cancer cell lines, whether also derived from lung carcinoma (A549-WT, HCC-827) or from other cancer types (cervical cancer for HeLa, pancreatic adenocarcinoma for MiaPaCa-2, triple-negative breast cancer for Hs578T). To test whether the increased cell lysis elicited by the selected variants stemmed from an intrinsically higher toxicity or simply from a more efficient replication, we compared the DNA replication speed of SVA, the most lytic variant, with HAdV-C5-WT. In A549-WT cells, the SVA variant replicated substantially faster than HAdV-C5-WT, both in terms of virus genome copies per cells and VP release in the culture supernatant (Figure 3c). Of note, the selection criteria was completion of full replication cycles rather than strictly transduction, which explains why variants hardly more infectious but strongly replicating and lytic were able to outcompete their competitors.

## DISCUSSION

Here we described Adenovirus directed evolution protocols facilitating the generation of highly complex libraries reaching close to one million variants (Figure 2a). This represents an approximately hundred-fold improvement compared with the most complex previously published AdV libraries (19). Although ADEVO libraries still fall short of the highest complexities reported with AAV vector libraries, they are likely to contain improved variants for numerous applications, provided that vector random modifications are performed in a relevant location and selection criteria are stringent enough to enrich the improved variants.

In addition, we focused here on user-friendly, relatively low-cost protocols requiring only materials readily available in most virology laboratories. Library complexity may be further increased with limited workflow changes, for example larger scale library generation or the use of specialized vector particles producer cell lines such as HEK293A or AD293 cells. Additional modifications could include the use of semi-random codons to decrease the proportion of non-functional variants with stop codons in the initial libraries.

We used random octopeptides (counting semi-random positions) inserted in the fiber knob HI loop because this type of inserts already facilitated efficient vector retargeting in various rational design studies (12–14). Therefore, less than 0.1% of the insert sequence space comprising several billion theoretical peptides (Figure 1c) was covered by our libraries. This means that there may exist more efficient inserts than the ones selected here: although the enriched variants overcame the selective pressure better than the other variants from the libraries, only a small fraction of the total pool of possible variants was generated and submitted to selection to begin with. However, there is to our knowledge no reason to assume that, using the same library generation protocols, library complexity would be higher with inserts of different length. Likewise, limiting insert length to four random amino acids in order to cover the full sequence space would not have guaranteed that more efficient variants could have been selected.

Round 0 libraries already showed extensive variations in abundance between variants (Figure 2b), even before selection had begun. This does not appear to stem from bias during variants genome cloning prior to transfection, given that the most prevalent variants in sequenced genome libraries did not represent more than 0.0008% of these libraries (Supplementary Table 2). The bias may instead result from random founder effects, whereby limited numbers of transfected vector genomes find excellent conditions to replicate in producer cells, while others suffer from suboptimal timing, intracellular or intranuclear location, cell health or cell cycle step. Furthermore, differential capsid stability at the vector particle packaging stage of the AdV life cycle may contribute to abundance discrepancies between variants. It is also possible that certain variants are able to undergo a second replication cycle before the vector library is harvested. This may explain that inserts with stop codon were already less frequent at round 0 than expected from purely random distribution, ranging from 4% to 12% (Figure 2d) compared to theoretical frequencies of 20% for Protocol 1 and 17% for Protocols 2 and 3. On the contrary, genome libraries matched the theoretical stop codon frequencies (Supplementary Table 2), confirming that genome library bias was very limited. Biases in round 0 library composition may be avoided by earlier VP harvest, which may increase library complexity at the cost of lower VP titers. This trade-off may explain the lower complexity of Protocols 2(b) and 3 libraries compared to Protocols 1 and 2(a) libraries, since the former displayed wider abundance distribution (Figure 2b), less inserts with stop codons (Figure 2d) and more extensive producer cell death (data not shown), suggesting that a noticeable proportion of the library variants already amplified through a second generation before round 0 VP library collection.

The lack of highly prevalent variant in genome libraries (Supplementary Table 2) also shows that the PCR amplification of inserts from library aliquots during NGS sequencing preparation did not cause substantial biases in detected insert prevalence. Likewise, previous studies reported that PCR amplifications prior to NGS variant calling caused only limited bias in reported library complexity (37,38). These articles recommended to use high input DNA mass, high-performance and high-fidelity polymerases and to maximise the percentage of PCR product sequenced, in order to decrease the bias in sequencing results. All these guidelines were satisfied by our workflows. Finally, the fact that several of the variants highly enriched after selection had given only a few reads or were not even detected by NGS in round 0 libraries (Figure 2f) indicates that many or most low-prevalence reads correspond to actual variants and not PCR or NGS errors, and that numerous variants were not included in the sequenced round 0 library aliquots. This suggested that library complexity may have been underestimated due to lack of sequencing depth, rather than overestimated through sequencing errors, and warranted the use of statistical modeling to estimate total library complexity. However, modeling appeared to lack robustness. For libraries where more than three library aliquots had been sequenced, we calculated the estimated complexity for all possible combinations of three aliquots and found up to three-fold variations depending primarily on the total number of reads per aliquot (Supplementary Figure 2). Furthermore, the estimates given by different relevant models also varied up to four-fold. Fortunately, the ratios between the estimated complexities of different libraries were always similar and roughly matched the ratios of observed complexities (i.e. the number of variants identified by NGS in the sequenced aliquots). We therefore assume that statistical models enable to compare the relative complexities of different libraries, although some of them may give inaccurate estimates. We ensured estimation consistency by always following the same protocols for NGS and analysis and focused on the model which gave the results closest to the average of all relevant models.

HAdV-C5-WT was able to infect A549-DCAR cells relatively efficiently (Figure 3b). Since no trace of residual CAR expression could be detected at the genomic (24) nor at the protein level (data not shown), this permissiveness was likely due to the strong expression by A549-derived cells of secondary adenovirus receptors including heparan sulfate proteoglycans and integrins (39,40). Therefore, the selective pressure on target cell infectivity was substantially lower than originally desired. This could provide at least a partial explanation why the most highly enriched variants did not differ strongly from HAdV-C5-WT based on infectivity and cell type specificity, but rather the selective pressure favored variants with improved replication and cell lysis kinetics. This could also explain why the selection process appeared to be slow, with no distinct insert sequence pattern having emerged at round 4 (Figure 2e), and extensive variations in variant prevalence still occurring in subsequent rounds (Figure 2f). It is likely that selection would have been faster if stronger selective pressures had been applied, for example using less permissive cells as target, fully detargeted backbones for library generation that do not bind known adenovirus secondary receptors, or anti-integrin antibodies or RGD peptides to saturate target cells integrins prior to library infection. Alternatively, if difficult selection settings require it, selection speed may be optimized by decreasing target cell multiplicity of infection, the time during which VPs are allowed to infect target cells before media change, or selection round duration.

The most highly enriched variants were re-cloned from assembly backbones after identification and purified individually. This ensures that the differences they displayed compared to HAdV-C5-WT during characterization (Figure 3) come from their fiber insert and not potential mutations elsewhere in the genome that may have arisen during library generation and selection. The oncolytic efficiency of the selected variants correlated with the electric charge of their insert peptide and with the presence of potential furin cleavage sites. Indeed, the FYK variant has 2 positively charged amino acids in its insert, CLY has 3, WSR, SRG and SVA have 5, and LRT 6. Meanwhile, the canonical RX(K/R)R furin cleavage site is present once in the SRG and SVA variants inserts and twice in the LRT variant insert, the WSR variant carries one RXXR minimal furin recognition motif, and the ProP software (41) predicted that the LRT insert indeed had a significant furin cleavage potential (Supplementary Figure 3). We therefore hypothesize that the potential fiber proteolytic cleavage or the fiber electric charge was responsible for the increased DNA replication and cell lysis. The fiber protein was not known to be involved in DNA replication, but the inserts may confer it new functions, for example the activation or inhibition of new ligands involved in a cellular signaling pathway respectively beneficial or detrimental for AdV replication. The inserts may also indirectly influence the replication process for example by modifying the kinetics and efficiency of vector particle endosomal escape and intracellular trafficking to the nucleus. Investigations of the mechanisms of fiber-mediated replication increase are warranted.

The lower cell lysis ability of the CLY, FYK and WSR variants compared to LRT, SRG and SVA (Figure 3d) may hint toward infection- and lysis-independent pathways through which fiber inserts could increase OAdV fitness. Alternatively, this can be interpreted as showing a lower fitness of the former variants compared to the latter, since CLY and FYK were selected from Protocol 1 library and did not have to compete with the other characterized variants, and WSR seemed to have a slower enrichment rate than its counterparts in Protocols 2 and 3 libraries (Figure 2f).

All three ADEVO protocols facilitated high complexity library generation. Protocols 2 and 3 libraries performed very similarly, even selecting to a large extent the same variants. The studied Protocol 1 library displayed higher purity than Protocols 2 and 3 libraries (Figure 2c) and appeared to reach relative equilibrium faster during selection than library a of Protocol 2. Indeed, highly enriched variants could already be identified at selection round 4 (Figure 2f) and the insert logo profile did not substantially change between rounds 4 and 8 (Figure 2e). Moreover, Protocols 2 and 3 libraries contained more bystander variants that had reached high prevalence already at selection round 0 (Figure 2b) apparently through random founder effects, suggesting that Protocol 1 provided a more equal starting point to all variants before selection. However, Protocol 1 required more time and resources due to the two-steps library construction. The variants selected by Protocol 1 performed worse than Protocols 2 and 3 enriched variants as oncolytic vectors, but this may be due to the insert flanking sequences or the lack of strong selective pressure for the intended goal of tropism retargeting rather than to differences in library generation protocol. The large overlap between variants that were highly enriched in either of the three Protocols 2 and 3 libraries indicate that the selection was robust, but also that round 0 libraries already shared numerous variants despite having been generated independently and covering only a limited fraction of the theoretical sequence space. Library complexity may thus have been limited by biases in the composition of random oligonucleotides used for library generation, although these potential biases were not extensive enough as to visibly skew the insert amino acid distribution in the whole library (Figure 2g).

Although ADEVO protocols proved satisfactory in library generation, this proof-of-concept study highlights the necessity of carefully designing stringent selection protocols for clinical applications. In particular, vector retargeting should rely on selection independent of vector replication and cell lysis. Suitable protocols would therefore not only be applicable to OAdVs, but also to replication-deficient gene delivery vectors. Furthermore, ADEVO may be harnessed for other goals than tropism retargeting, for example escape from neutralizing antibodies and other serum proteins, whose binding to the AdV capsid can cause therapeutic inefficiency or even vector toxicity (42–44). Detargeting from these undesired ligands could be achieved by random peptide insertion in other fiber knob sites than the HI loops, as well as in other capsid proteins, notably in hexon hyper variable regions which are the main binding site of undesired ligands (45,46). Finally, other types than HAdV-C5 may be used as backbones in order to combine specific improvements obtained by directed evolution with the natural diversity in potentially clinically interesting AdV features.

In this study, we established user-friendly AdV directed evolution workflows covering random library generation, selection and analysis. We achieved unprecedented library complexity for this type of vector and were able to select variants with increased oncolytic potential. We hope that this study provides the necessary toolkit for a wider use of directed evolution in the generation of novel Adenovirus vectors for a broad range of applications. Furthermore, the fiber proteins selected in this study may be utilized to construct improved oncolytic vectors, provided that the potential safety concern raised by their lack of cancer type specificity be addressed by appropriate approaches such as cancer-specific promoters and conditional replication.

## DATA AVAILABILITY

The main python codes used in this study to characterize NGS-sequenced libraries are available on zenodo in open access (DOI: 10.5281/zenodo.10083525). All other data necessary for the analysis and reproduction of the results presented in the article is available in the article and supplements or will be made available by the authors upon reasonable request. Supplementary Data are available at NAR online.

## ACKNOWLEDGEMENTS

We are grateful to Zhi Hong Lu (Washington University School of Medicine) for the insightful discussion on potential pathways of fiber inserts gain-of-function.

## REFERENCES

1. Davison, A.J., Benko, M. and Harrach, B. (2003) Genetic content and evolution of adenoviruses. J Gen Virol, 84, 2895–2908.

2. Human Adenovirus Working Group. (2023). http://hadvwg.gmu.edu/

3. Sakurai, F., Tachibana, M. and Mizuguchi, H. (2022) Adenovirus vector-based vaccine for infectious diseases. Drug Metab Pharmacokinet, 42, 100432.

4. Gene Therapy Clinical Trials Worldwide. (2023). The Journal of Gene Medicine. https://a873679.fmphost.com/fmi/webd/GTCT

5. Zhao, Z., Anselmo, A.C. and Mitragotri, S. (2022) Viral vector-based gene therapies in the clinic. Bioeng Transl Med, 7, e10258.

6. Eissa, I.R., Bustos-Villalobos, I., Ichinose, T., Matsumura, S., Naoe, Y., Miyajima, N., Morimoto, D., Mukoyama, N., Zhiwen, W., Tanaka, M. et al. (2018) The Current Status and Future Prospects of Oncolytic Viruses in Clinical Trials against Melanoma, Glioma, Pancreatic, and Breast Cancers. Cancers (Basel), 10.

7. Liang, M. (2018) Oncorine, the World First Oncolytic Virus Medicine and its Update in China. Curr Cancer Drug Targets, 18, 171–176.

8. Gao, J., Zhang, W. and Ehrhardt, A. (2020) Expanding the Spectrum of Adenoviral Vectors for Cancer Therapy. Cancers (Basel), 12.

9. Yamamoto, Y., Hiraoka, N., Goto, N., Rin, Y., Miura, K., Narumi, K., Uchida, H., Tagawa, M. and Aoki, K. (2014) A targeting ligand enhances infectivity and cytotoxicity of an oncolytic adenovirus in human pancreatic cancer tissues. J Control Release, 192, 284–293.

10. Wesseling, J.G., Bosma, P.J., Krasnykh, V., Kashentseva, E.A., Blackwell, J.L., Reynolds, P.N., Li, H., Parameshwar, M., Vickers, S.M., Jaffee, E.M. et al. (2001) Improved gene transfer efficiency to primary and established human pancreatic carcinoma target cells via epidermal growth factor receptor and integrin-targeted adenoviral vectors. Gene Ther, 8, 969–976.

11. Owczarek, C., Elmasry, Y. and Parsons, M. (2023) Contributions of coxsackievirus adenovirus receptor to tumorigenesis. Biochem Soc Trans, 51, 1143–1155.

12. Krasnykh, V., Dmitriev, I., Mikheeva, G., Miller, C.R., Belousova, N. and Curiel, D.T. (1998) Characterization of an adenovirus vector containing a heterologous peptide epitope in the HI loop of the fiber knob. J Virol, 72, 1844–1852.

13. Kritz, A.B., Nicol, C.G., Dishart, K.L., Nelson, R., Holbeck, S., Von Seggern, D.J., Work, L.M., McVey, J.H., Nicklin, S.A. and Baker, A.H. (2007) Adenovirus 5 fibers mutated at the putative HSPG-binding site show restricted retargeting with targeting peptides in the HI loop. Mol Ther, 15, 741–749.

14. Kurachi, S., Koizumi, N., Sakurai, F., Kawabata, K., Sakurai, H., Nakagawa, S., Hayakawa, T. and Mizuguchi, H. (2007) Characterization of capsid-modified adenovirus vectors containing heterologous peptides in the fiber knob, protein IX, or hexon. Gene Ther, 14, 266–274.

15. Wang, D., Tai, P.W.L. and Gao, G. (2019) Adeno-associated virus vector as a platform for gene therapy delivery. Nat Rev Drug Discov, 18, 358–378.

16. Kuhn, I., Harden, P., Bauzon, M., Chartier, C., Nye, J., Thorne, S., Reid, T., Ni, S., Lieber, A., Fisher, K. et al. (2008) Directed evolution generates a novel oncolytic virus for the treatment of colon cancer. PLoS One, 3, e2409.

17. Moreno, V., Barretina-Ginesta, M.P., Garcia-Donas, J., Jayson, G.C., Roxburgh, P., Vazquez, R.M., Michael, A., Anton-Torres, A., Brown, R., Krige, D. et al. (2021) Safety and efficacy of the tumor-selective adenovirus enadenotucirev with or without paclitaxel in platinum-resistant ovarian cancer: a phase 1 clinical trial. J Immunother Cancer, 9.

18. Dalkara, D., Byrne, L.C., Klimczak, R.R., Visel, M., Yin, L., Merigan, W.H., Flannery, J.G. and Schaffer, D.V. (2013) In vivo-directed evolution of a new adeno-associated virus for therapeutic outer retinal gene delivery from the vitreous. Sci Transl Med, 5, 189ra176.

19. Yamamoto, Y., Goto, N., Miura, K., Narumi, K., Ohnami, S., Uchida, H., Miura, Y., Yamamoto, M. and Aoki, K. (2014) Development of a novel efficient method to construct an adenovirus library displaying random peptides on the fiber knob. Mol Pharm, 11, 1069–1074.

20. Tabebordbar, M., Lagerborg, K.A., Stanton, A., King, E.M., Ye, S., Tellez, L., Krunnfusz, A., Tavakoli, S., Widrick, J.J., Messemer, K.A. et al. (2021) Directed evolution of a family of AAV capsid variants enabling potent muscle-directed gene delivery across species. Cell, 184, 4919–4938 e4922.

21. Nishimoto, T., Yoshida, K., Miura, Y., Kobayashi, A., Hara, H., Ohnami, S., Kurisu, K., Yoshida, T. and Aoki, K. (2009) Oncolytic virus therapy for pancreatic cancer using the adenovirus library displaying random peptides on the fiber knob. Gene Ther, 16, 669–680.

22. Nishimoto, T., Yamamoto, Y., Yoshida, K., Goto, N., Ohnami, S. and Aoki, K. (2012) Development of peritoneal tumor-targeting vector by in vivo screening with a random peptide-displaying adenovirus library. PLoS One, 7, e45550.

23. Miciak, J.J., Hirshberg, J. and Bunz, F. (2018) Seamless assembly of recombinant adenoviral genomes from high-copy plasmids. PLoS One, 13, e0199563.

24. Tsoukas, R.L., Volkwein, W., Gao, J., Schiwon, M., Bahlmann, N., Dittmar, T., Hagedorn, C., Ehrke-Schulz, E., Zhang, W., Baiker, A. et al. (2022) A Human In Vitro Model to Study Adenoviral Receptors and Virus Cell Interactions. Cells, 11.

25. Zhang, W., Fu, J., Liu, J., Wang, H., Schiwon, M., Janz, S., Schaffarczyk, L., von der Goltz, L., Ehrke-Schulz, E., Dorner, J., et al. (2017) An Engineered Virus Library as a Resource for the Spectrum-wide Exploration of Virus and Vector Diversity. Cell Rep, 19, 1698–1709.

26. Schroer, K., Arakrak, F., Bremke, A., Ehrhardt, A. and Zhang, W. (2022) HEHR: Homing Endonuclease-Mediated Homologous Recombination for Efficient Adenovirus Genome Engineering. Genes (Basel), 13.

27. Riedl, A., Fischer, J., Burgert, H.G. and Ruzsics, Z. (2022) Rescue of Recombinant Adenoviruses by CRISPR/Cas-Mediated in vivo Terminal Resolution. Front Microbiol, 13, 854690.

28. Jager, L., Hausl, M.A., Rauschhuber, C., Wolf, N.M., Kay, M.A. and Ehrhardt, A. (2009) A rapid protocol for construction and production of high-capacity adenoviral vectors. Nat Protoc, 4, 547–564.

29. Westhaus, A., Cabanes-Creus, M., Jonker, T., Sallard, E., Navarro, R.G., Zhu, E., Baltazar Torres, G., Lee, S., Wilmott, P., Gonzalez-Cordero, A. et al. (2022) AAV-p40 Bioengineering Platform for Variant Selection Based on Transgene Expression. Hum Gene Ther, 33, 664–682.

30. McKinney. (2010) Data structures for statistical computing in python. Proceedings of the 9th Python in Science Conference, 445, 56–61.

31. Harris, C.R., Millman, K.J., van der Walt, S.J., Gommers, R., Virtanen, P., Cournapeau, D., Wieser, E., Taylor, J., Berg, S., Smith, N.J., et al. (2020) Array programming with NumPy. Nature, 585, 357–362.

32. Cock, P.J., Antao, T., Chang, J.T., Chapman, B.A., Cox, C.J., Dalke, A., Friedberg, I., Hamelryck, T., Kauff, F., Wilczynski, B. et al. (2009) Biopython: freely available Python tools for computational molecular biology and bioinformatics. Bioinformatics, 25, 1422–1423.

33. Hunter, J.D. (2007) Matplotlib: A 2D Graphics Environment. Computing in Science & Engineering, 9, 90–95.

34. Chao, A. (1987) Estimating the Population Size for Capture-Recapture Data with Unequal Catchability. Biometrics, 43.

35. Schmit, P.F., Pacouret, S., Zinn, E., Telford, E., Nicolaou, F., Broucque, F., Andres-Mateos, E., Xiao, R., Penaud-Budloo, M., Bouzelha, M. et al. (2020) Cross-Packaging and Capsid Mosaic Formation in Multiplexed AAV Libraries. Mol Ther Methods Clin Dev, 17, 107–121.

36. Truong, C.S. and Yoo, S.Y. (2022) Oncolytic Vaccinia Virus in Lung Cancer Vaccines. Vaccines (Basel), 10.

37. Rochette, N.C., Rivera-Colón, A.G., Walsh, J., Sanger, T.J., Campbell-Staton, S.C. and Catchen, J.M. (2022) On the causes, consequences, and avoidance of PCR duplicates: towards a theory of library complexity. BioRxiv.

38. Ebbert, M.T., Wadsworth, M.E., Staley, L.A., Hoyt, K.L., Pickett, B., Miller, J., Duce, J., Alzheimer’s Disease Neuroimaging, I., Kauwe, J.S. and Ridge, P.G. (2016) Evaluating the necessity of PCR duplicate removal from next-generation sequencing data and a comparison of approaches. BMC Bioinformatics, 17 Suppl 7, 239.

39. Davison, E., Kirby, I., Whitehouse, J., Hart, I., Marshall, J.F. and Santis, G. (2001) Adenovirus type 5 uptake by lung adenocarcinoma cells in culture correlates with HAdV-C5 fibre binding is mediated by alpha(v)beta1 integrin and can be modulated by changes in beta1 integrin function. J Gene Med, 3, 550–559.

40. Dechecchi, M.C., Melotti, P., Bonizzato, A., Santacatterina, M., Chilosi, M. and Cabrini, G. (2001) Heparan sulfate glycosaminoglycans are receptors sufficient to mediate the initial binding of adenovirus types 2 and 5. J Virol, 75, 8772–8780.

41. Duckert, P., Brunak, S. and Blom, N. (2004) Prediction of proprotein convertase cleavage sites. Protein Eng Des Sel, 17, 107–112.

42. Tian, J., Xu, Z., Moitra, R., Palmer, D.J., Ng, P. and Byrnes, A.P. (2022) Binding of adenovirus species C hexon to prothrombin and the influence of hexon on vector properties in vitro and in vivo. PLoS Pathog, 18, e1010859.

43. Baker, A.T., Boyd, R.J., Sarkar, D., Teijeira-Crespo, A., Chan, C.K., Bates, E., Waraich, K., Vant, J., Wilson, E., Truong, C.D. et al. (2021) ChAdOx1 interacts with CAR and PF4 with implications for thrombosis with thrombocytopenia syndrome. Sci Adv, 7, eabl8213.

44. Waddington, S.N., McVey, J.H., Bhella, D., Parker, A.L., Barker, K., Atoda, H., Pink, R., Buckley, S.M., Greig, J.A., Denby, L. et al. (2008) Adenovirus serotype 5 hexon mediates liver gene transfer. Cell, 132, 397–409.

45. Atasheva, S., Emerson, C.C., Yao, J., Young, C., Stewart, P.L. and Shayakhmetov, D.M. (2020) Systemic cancer therapy with engineered adenovirus that evades innate immunity. Sci Transl Med, 12.

46. Alba, R., Bradshaw, A.C., Parker, A.L., Bhella, D., Waddington, S.N., Nicklin, S.A., van Rooijen, N., Custers, J., Goudsmit, J., Barouch, D.H., et al. (2009) Identification of coagulation factor (F)X binding sites on the adenovirus serotype 5 hexon: effect of mutagenesis on FX interactions and gene transfer. Blood, 114, 965–971.

