## Supplementary figures and images for "ADEVO: Proof-of-concept of Adenovirus Directed EVOlution by random peptide display on the fiber knob"

### Supplementary Figure 1

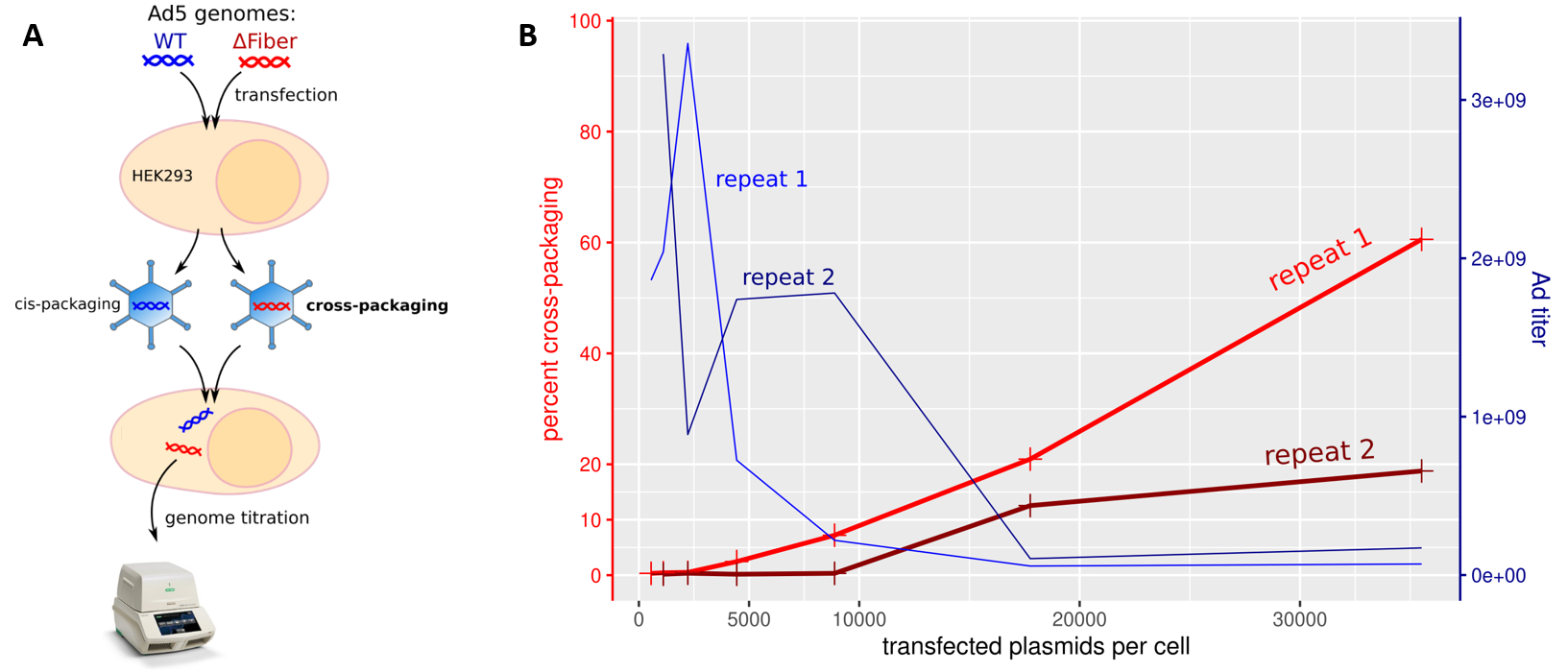

### Supplementary Figure 2

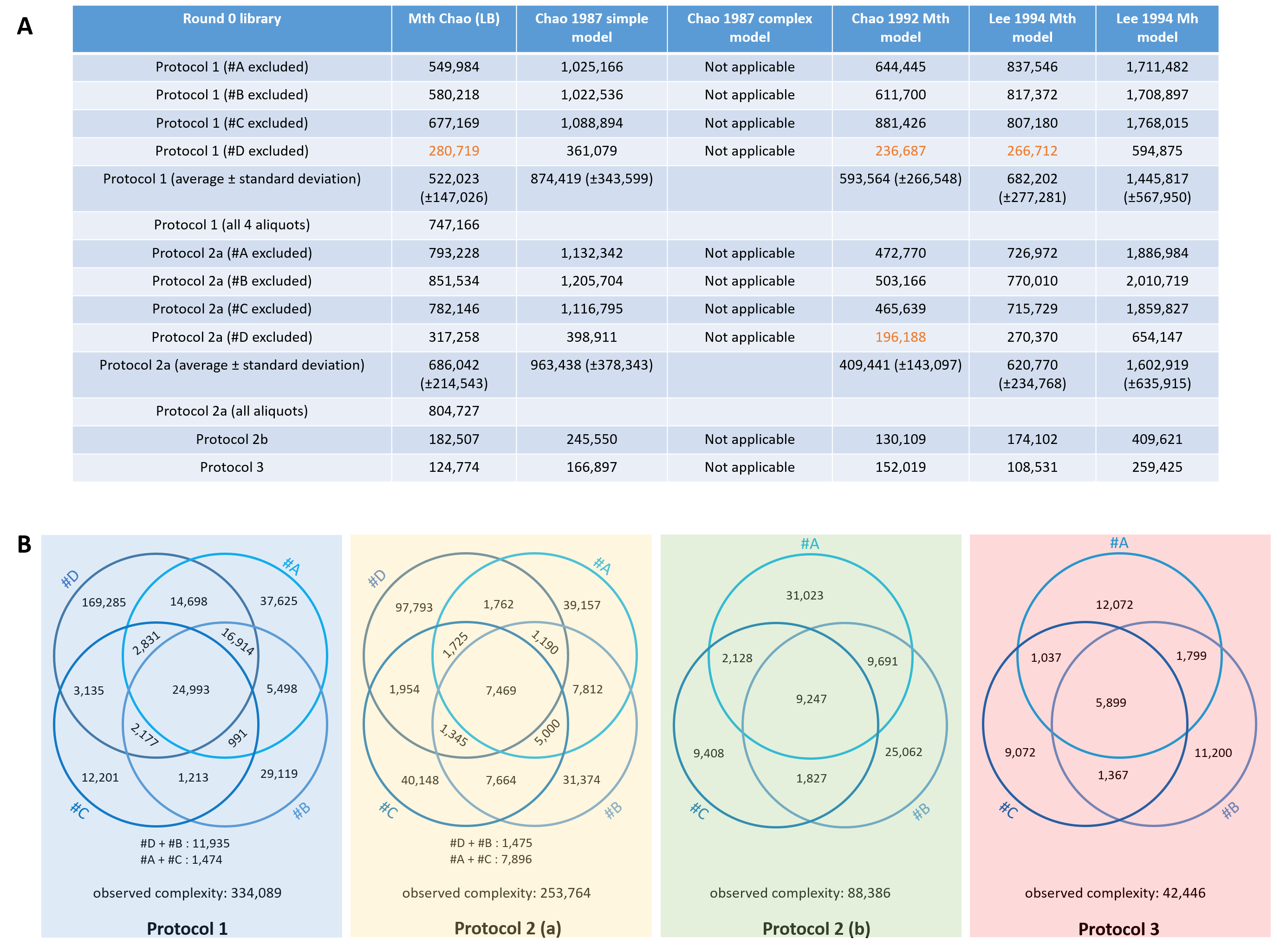

### Supplementary Figure 3

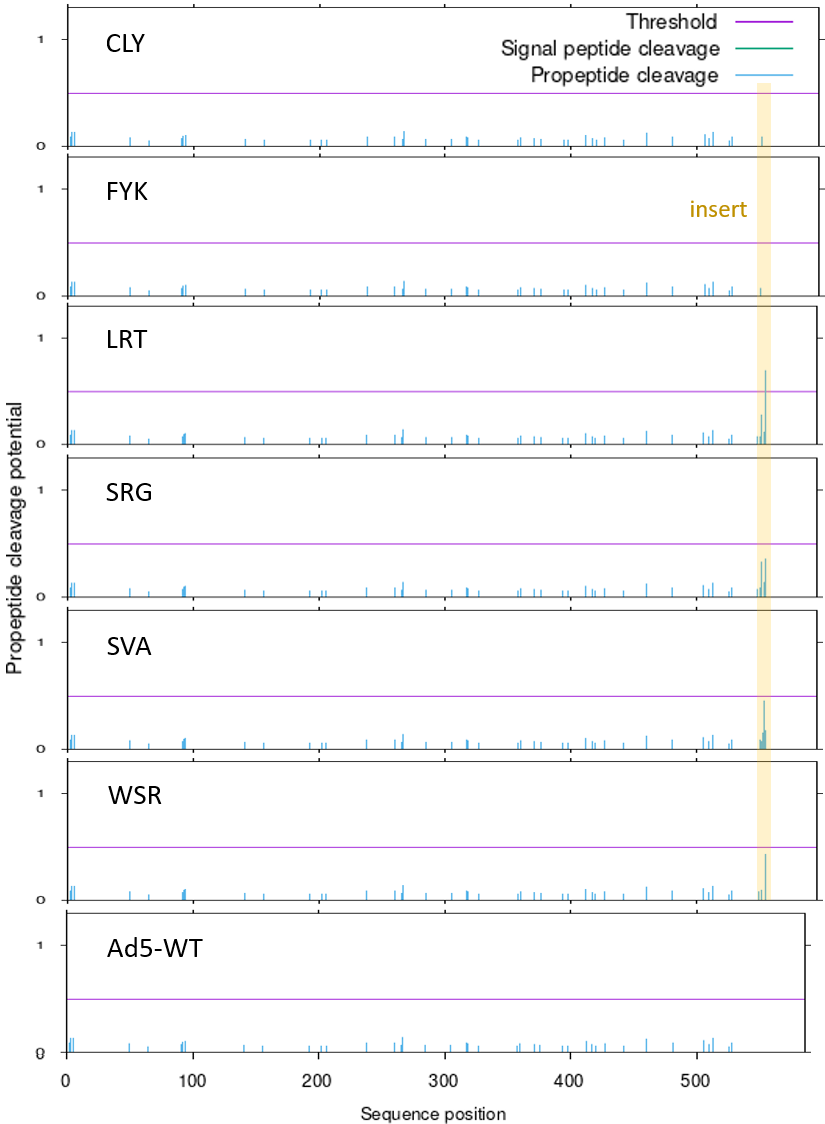

### Supplementary Figure 4

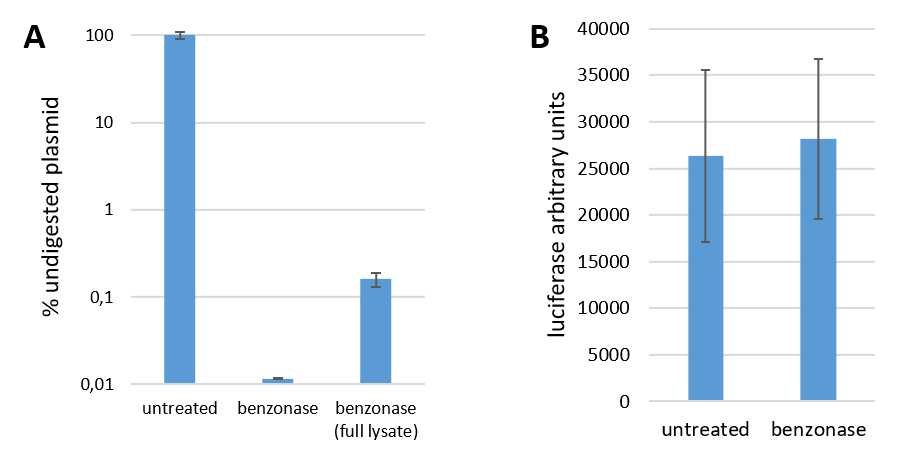
